# ClusterV-Web: A User-Friendly Tool for Profiling HIV Quasispecies and Generating Drug Resistance Reports from Nanopore Long-Read Data

**DOI:** 10.1101/2023.10.19.563033

**Authors:** Junhao Su, Shumin Li, Zhenxian Zheng, Tak-Wah Lam, Ruibang Luo

**Affiliations:** University of Hong Kong

## Abstract

**Summary:** Third-generation long-read sequencing is an increasingly utilized technique for profiling HIV quasispecies and detecting drug resistance mutations due to its ability to cover the entire viral genome in individual reads. Recently, the ClusterV tool has demonstrated accurate detection of HIV quasispecies from Nanopore long-read sequencing data. However, the need for scripting skills and a computational environment may act as a barrier for many potential users. To address this issue, we have introduced ClusterV-Web, a user-friendly web-based application that enables easy configuration and execution of ClusterV, both remotely and locally. Our tool provides interactive tables and data visualizations to aid in the interpretation of results. This development is expected to democratize access to long-read sequencing data analysis, enabling a wider range of researchers and clinicians to efficiently profile HIV quasispecies and detect drug resistance mutations.

**Availability and implementation:** ClusterV-Web is freely available and open source, with detailed documentation accessible at http://www.bio8.cs.hku.hk/ClusterVW/. The standalone Docker image and source code are also available at https://github.com/HKU-BAL/ClusterV-Web.

**Contact:**, Department of Computer Science, The University of Hong Kong, Hong Kong, China, Department of Computer Science, The University of Hong Kong, Hong Kong, China

**Supplementary information:** None

## 1 Introduction

The human immunodeficiency virus (HIV) demonstrates extensive genetic heterogeneity both between and within hosts, owing to its high mutation and recombination rate. Drug-resistance mutations become a major barrier to clinical treatment (Yeo, et al., 2020). Therefore, closely monitoring HIV quasispecies (distinct, closely related viral mutants) is critical for controlling HIV infection and studying the virus-host coevolution mechanism. High-throughput sequencing-based methods has recently emerged for profiling HIV quasispecies. However, even with low sequencing error, short-read sequencing struggles to reconstruct haplotypes due to uneven depth and high viral genome similarities. In this case, long-read sequencing technologies, such as Oxford Nanopore Technologies (ONT) and PacBio Single-Molecule Real-Time (SMRT) sequencing, have become a promising option as their long read can cover the entire viral genomes (Jiao, et al., 2022). ONT, in particular, could be a popular option in clinical diagnosis due to its portability, speed, and low cost. The main difficulty of applying long-read sequencing to HIV quasispecies discovery is their relatively high base error. Several novel analysis methods have been proposed for long-read sequencing and targeted to address the data analysis with the high error rates. For instance, QuasiSeq produces high-quality quasispecies sequences with PacBio long-read data through signature-based clustering (Jiao, et al., 2022). Strainline can assemble strain-resolved viral quasispecies without a reference genome(Luo, et al., 2022), while CliqueSNV is reference-based method for haplotype reconstruction based on the linkage between single nucleotide variations (SNVs)(Knyazev, et al., 2021).

Previously, we proposed ClusterV, a pipeline for accurately detecting HIV quasispecies from ONT data and identifying the HIV drug-resistance mutations (Ng, et al., 2023). ClusterV has demonstrated high sensitivity and precision in 59 HIV-positive plasma samples and has been cross-validated with short-read as well as Sanger sequencing. However, ClusterV was released as a command-line tool that assumed potential users had command-line skills and could set up the dependency environment, which could limit its accessibility to a broader user base. Additionally, various types of output files were generated, requiring additional effort to integrate them before interpretation.

To address these limitations, we developed ClusterV-Web. This tool wraps ClusterV into an interactive web server that offers a user-friendly interface for running tasks, and auto-generates result reports and data visualizations using our computing nodes. ClusterV-Web is freely accessible at http://www.bio8.cs.hku.hk/ClusterVW/. We also provide a standalone docker image to run ClusterV-Web locally at https://hub.docker.com/r/hkubal/clustervw. We expect ClusterV-Web will facilitate the identification process of HIV quasispecies and make the analysis feasible for a wide range of audiences.

## 2 Results

### 2.1 Software implementation

ClusterV-Web is a web-based application designed for the discovery of HIV quasispecies with or without drug-resistance, utilizing the ClusterV algorithm. We have deployed the online version of ClusterV-Web on a server equipped with 6 Intel i7 CPUs and 64GB of memory. In addition, we have created a standalone version of ClusterV-Web as a docker image that can be easily launched on machines with common CPU architectures (x86_64 and arm64) and mainstream operating systems (Windows, MacOS, and Linux distributions). The implementation is shown in Fig. 1A. Bootstrap 3, CSS, and Javascript are used to support the interactive user interface. We utilized Flask as the web framework and employed a SQLite database to manage tasks. A Redis server was launched within a computing cluster to execute the ClusterV processes. To submit background jobs and monitor their progress, we implemented a task queue using the Python RQ package. Compatibility testing of ClusterV-Web has been performed on various popular browsers, including Google Chrome, Apple Safari, and Mozilla Firefox.

**Fig. 1.**
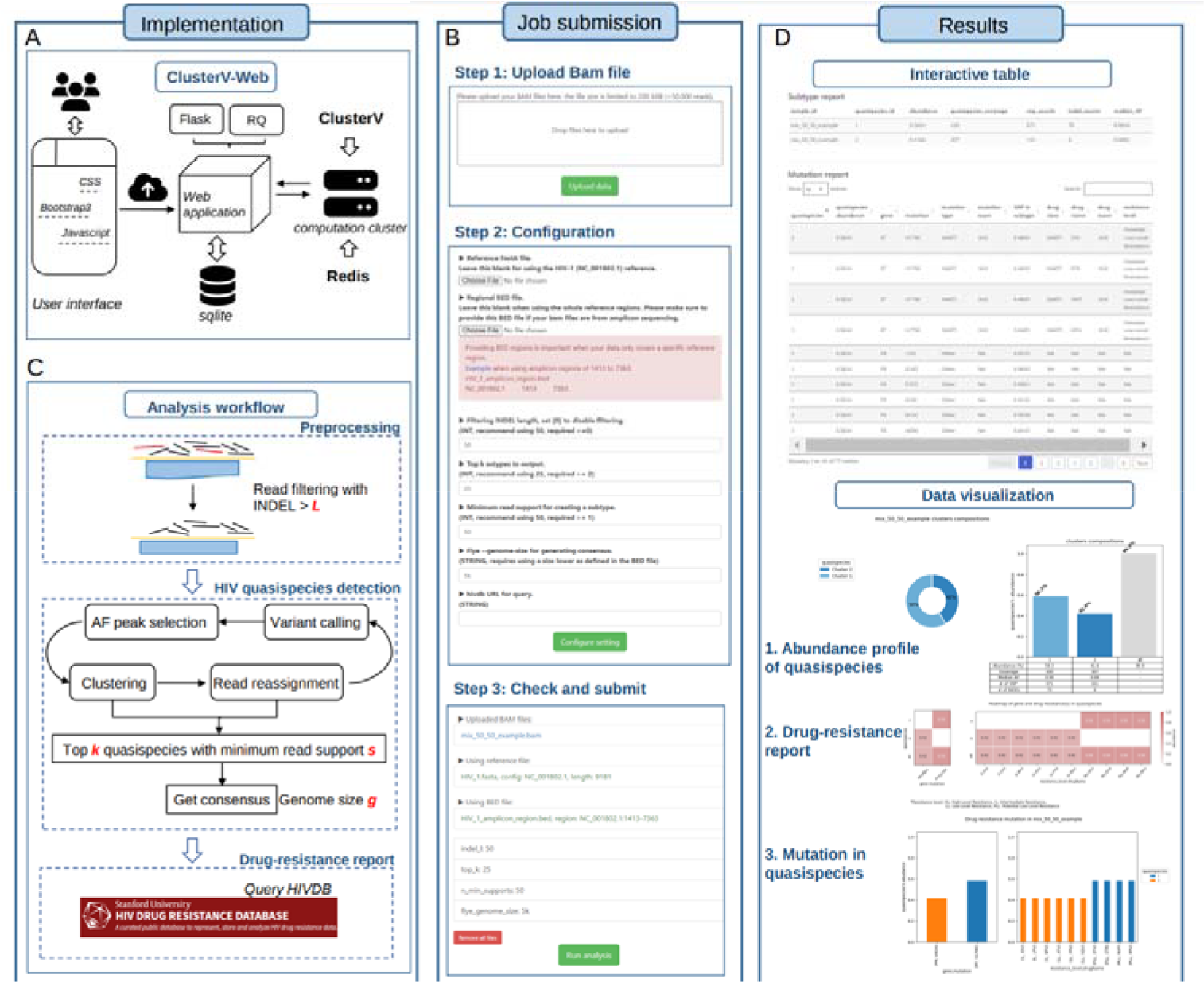
Overview and graphical interface of the ClusterV-Web. (**A**) Implementation of the web-application. (**B**) Steps of configuring and running a job. (**C**) Analysis pipeline with adjustable parameters. (**D**) Example results of interactive tables and data visualizations.

### 2.2 Workflow and usage

ClusterV-Web utilizes the state-of-the-art ClusterV pipeline for HIV quasispecies discovery and drug-resistance identification from ONT sequencing data. The online version is freely accessible without the need for installation, while the standalone version can be downloaded and launched using the following command:

#### docker run --name clustervw -d -p 8000:5000 --rm hkubal/clustervw:latest

As shown in Fig. 1B, the analysis with ClusterV-Web can be completed in three simple steps: 1) upload the input file, 2) configure the ClusterV pipeline parameters, and 3) check the settings and “Run analysis”. To ensure compatibility with various protocols, ClusterV-Web accepts reference-aligned reads in the BAM format as input. Users should prepare BAM files following best practices based on their device. The online version limits the BAM file size to 200MB, and users should switch to the limitless standalone local version if the file size exceeds this limit. Upon clicking “Upload data”, the data is anonymously sent to the server, and the configuration section is then presented.

In the configuration step, users can upload their customized reference genome in FASTA format if they prefer not to use HIV-1 (NC001802.1). If amplicon-based sequencing was used, providing the reference region BED file is highly recommended as it improves performance.

The best practice parameters for ClusterV are built into the ClusterV-Web interface, and users can adjust these parameters as necessary. The tunable parameters are for the three main modules of ClusterV (Fig. 1C). The first is preprocessing, with the filtering INDEL length parameter ***L*** set to 50 by default. Reads with large INDELs will be discarded. The second module is HIV quasispecies detection, which employs an iteratively read-based hierarchical clustering using variants called by Clair-ensemble (Leung, et al., 2022). Users may set the top ***k*** output quasispecies and minimum read support ***s*** of quasispecies to guide the algorithm.

The last module generates a drug-resistance report by querying the Stanford HIV Drug Resistance Database (HIVDB) (Shafer, 2006), and users may set their preferred genome size ***g*** of the built querying consensus (default: 5k) (Kolmogorov, et al., 2019).

All settings can be reviewed and submitted in step 3. ClusterV-Web employs an asynchronous job queue system that connects the running processes and the web application. The job status can be easily monitored through the progress bar on the results page. Once the job is completed, ClusterV-Web presents interactive tables that include detected HIV quasispecies and mutation details. Various data visualizations are provided to facilitate result interpretation (Fig1. D). Additionally, users can submit multiple tasks and manage them on the results page.

## 3 Conclusion

ClusterV-Web is an interactive visual analytical online tool for detecting and reporting drug-resistant HIV quasispecies from ONT data. The tool offers several key advantages to facilitate the analysis process. First, users can perform an analysis pipeline with minimal bioinformatics experience. Second, the installation of dependency for command-line tools can be tedious and prone to conflicts across different platforms. To address this issue, the online version of ClusterV-Web requires no installation, as the process runs on our computation cluster. Lastly, ClusterV-Web provides multiple interactive tables and figures that contain extensive information, such as detected quasispecies profiles and details of mutations. These features make it easy for users to combine and interpret results.

Third-generation long-read sequencing has emerged and proven to be highly effective in HIV quasispecies profiling, offering great potential for improving clinical treatment. Equipped with the high-performance ClusterV pipeline, the easy-to-use ClusterV-Web application is expected to significantly facilitate the analysis process.

## Acknowledgements

Not applicable.

## Supplementary data

Not applicable.

